# Development of Severity Index for Oral Epithelial Dysplasia Using Fuzzy Group Decision Making Algorithm

**DOI:** 10.1101/2025.01.20.633806

**Authors:** Debaleena Nawn, Pooja Lahiri, Budhaditya Dey, Mousumi Pal, Ranjan Rashmi Paul, Debashree Guha Adhya, Debjani Chakraborty, Jyotirmoy Chatterjee

**Affiliations:** Adamas University, Kolkata, India; Indian Institute of Technology, Kharagpur, India; R. Ahmed Dental College and Hospital, Kolkata, India; Gurunanak Institute of Dental Science and Research, Kolkata, India; JIS Institute of Advanced Studies and Research, Kolkata, India

**Keywords:** oral epithelial dysplasia, histopathological features, features’, weights, severity index, clinical decision support system, group decision making, fuzzy logic

## Abstract

**Background:** Oral epithelial dysplasia (OED) grading suffers from several levels of uncertainties and imprecision. An index is preferred in this context to improve reliability and reduce subjectivity in diagnostic decision making. In this study, a fuzzy logic-based disease severity index (SI) is formulated considering standard histopathological features used in OED grading.

**Methods:** Oral onco-pathologists were asked to independently provide weights of different features, according to their clinical significance in the context of dysplasia. Aggregated weight of each feature was calculated from individual assessment of different onco-pathologists by a fuzzy logic-based group decision making algorithm. Confidence levels of experts were also included to improve robustness of the method. SI was generated by integrating abnormality score of each feature with its weight. Abnormality degree of each feature was expressed in linguistic terms by onco-pathologists which were subsequently represented by a triangular fuzzy number. Fuzzy membership function was used as it can capture the ambiguity of experts’ opinion regarding abnormality of individual feature. Finally, defuzzification was used to get a crisp index from weighted sum of all features.

**Result:** SI was found to be statistically different (p<0.01) for different grades of dysplasia i.e mild, moderate and severe with added advantage of stratifying each grade in low and high subcategory.

**Conclusion:** Key contribution of our work is that we have developed a fuzzy logic-based group consensus process regarding weights of histopathological features for OED grading. Present methodology of developing SI can be applied to other medical decision-making problems as well.

**Highlights:**

- A severity index is proposed to improve reproducibility of oral epithelial dysplasia grading.
- Clinical significance of each histopathological and cytopathological feature in the context of dysplasia is reflected by its weight.
- A fuzzy logic-based group decision making algorithm is used to reduce subjectivity of features’ weights.
- Confidence level of oral onco-pathologists are given due importance in derivation of aggregated weight of each feature.
- Abnormality score of each feature is evaluated in fuzzy scale to capture clinicians’ ambiguity.
- Present methodology may be useful for developing indices of other diseases.

Graphical abstract

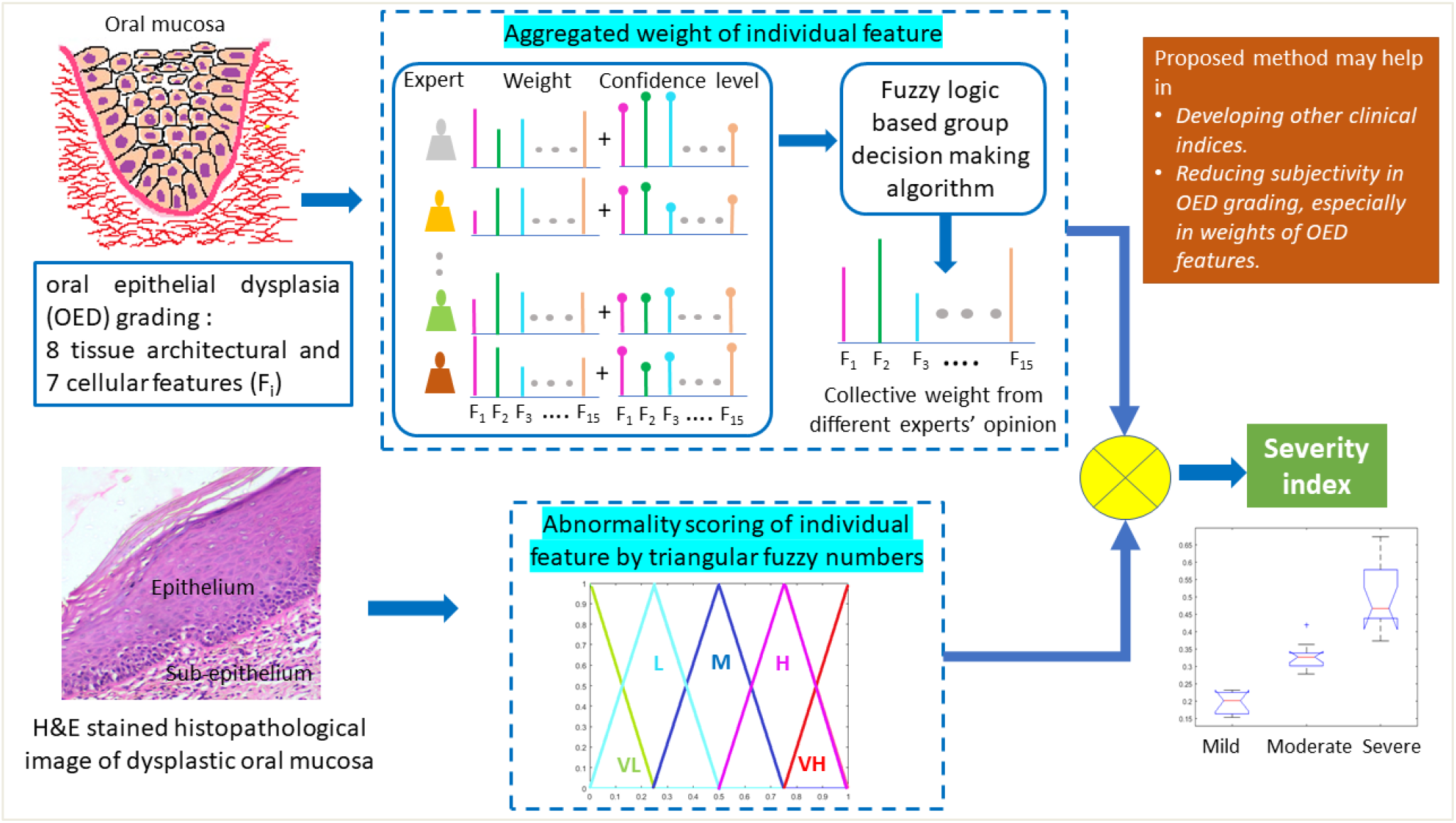

## 1. Introduction

Oral cancer (OC), a significant health problem worldwide, usually develops from pre-existing oral potentially malignant disorders (OPMDs) like oral sub-mucous fibrosis (OSF), leukoplakia (OLKP), erythroplakia [1]. OPMDs may or may not be associated with dysplasia while presence of dysplasia increases malignant potentiality of OPMDs [2, 3]. Oral epithelial dysplasia (OED) is progressively graded as mild, moderate and severe. Proper assessment of OED severity is very important for better therapeutic intervention and management which eventually enhances survival rate of patients. Among various grading systems of OED (viz. Smith and Pindborg photographic methods, Ljubljana classification, Squamous Intraepithelial Neoplasia/dysplasia classification), WHO classification system is most extensively followed [4, 5]. In all the methods, criteria for grading of OED are combination of both architectural and cytological alterations of surface epithelium.

Inter and intra observer variability is a well acknowledged concern while evaluation of OED severity [4]. In this context, an index may be preferred to report severity in a compact and consistent manner. In this study, we have formulated a disease severity index (SI) based on weight and degree of abnormality of each feature, to minimize decision making errors in OED assessment. We have consulted oral onco-pathologists of varying experience and each of them separately provided weightage or “weights” to the histological and cytological features. Weight implies significance of each feature in the context of OED. To minimize the subjectivity in weights, we attained consensus by a fuzzy logic-based group decision making problem which also considers confidence level of onco-pathologists regarding the weights. Previously, an epithelial atypia index for OED was proposed by Smith and Pindborg, where light microscopic histopathological features were given a weighted score based on their abnormality. However, the weightages given to each feature were highly subjective and not suitable for routine assessment. In SI calculation, abnormality score of each feature denotes degree of alteration of the feature from normal condition. While evaluating abnormality of each feature, pathologists find it difficult to grade the dysplastic features with exact numerical value due to its biological complexity and qualitative estimation. Linguistic assessment is preferred in such scenarios where complex assessment-based judgment is required. Several reports have shown suitability of fuzzy logic in medical decision making as it can encapsulate the vagueness and uncertainty of subjective evaluation [6-13]. Hence, experts have expressed the abnormality of each histopathological feature as a linguistic variable which are then transformed to fuzzy scale to achieve the SI of the disease.

## 2. Materials and methods

### 2.1 Inclusion of study samples and microscopic imaging

Formalin fixed and paraffin embedded oral biopsy specimens clinically representing precancerous lesions in oral cavity were obtained from Guru Nanak Institute of Dental Science and Research (GNIDSR), Kolkata, India. The biopsy was done from the buccal mucosa of patients and co-morbid samples were excluded. Histopathological analysis for the assessment of the disease state was performed on 4μm thick tissue sections using Haematoxylin and Eosin (H&E) staining. Microscopic images having size 1388 × 1040 pixels were grabbed digitally using bright-field inverted optical microscope (Zeiss Observer.Z1, Carl Zeiss, Germany) associated with CCD camera (AxioCam MRC, Carl Zeiss) under 20x and 40x objective (Fig 1).

**Fig. 1.**
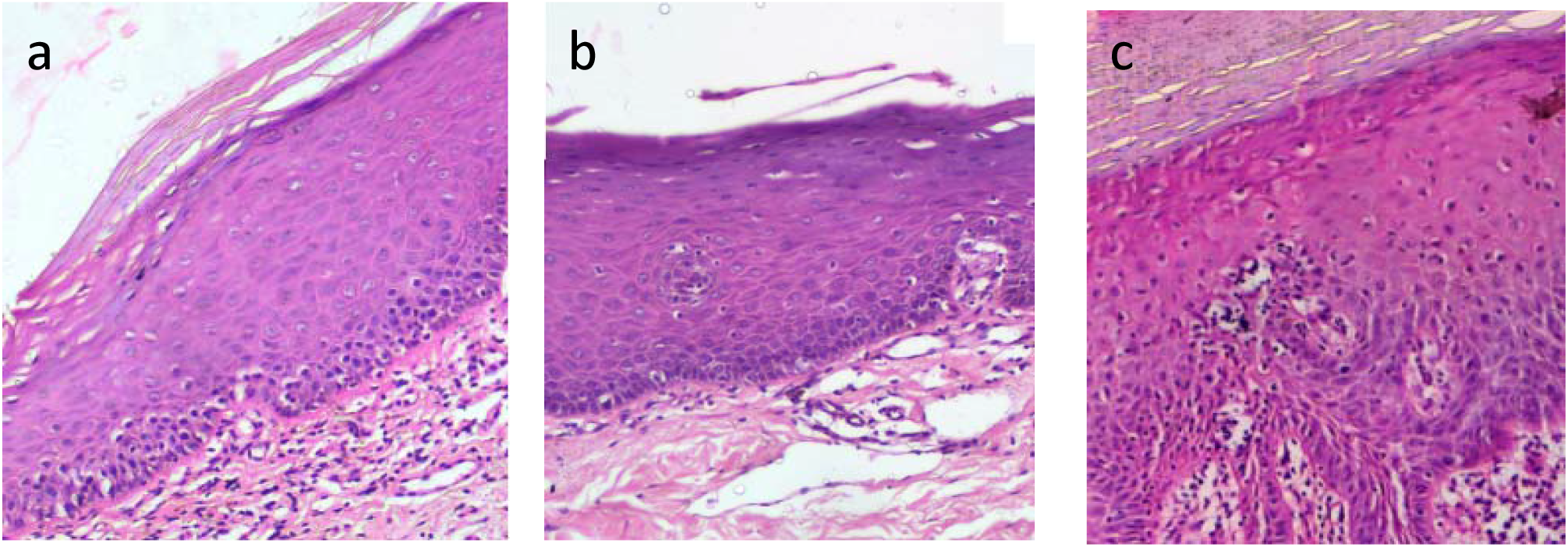
Light microscopic H&E stained histopathological images (200×) of dysplastic oral mucosa (a) mild (b) moderate and (c) severe.

#### Ethical statement

This study was performed with the informed consent of all individuals and approved by institutional ethics committee (GNIDSR/IEC/07/16).

### 2.2 Oral epithelial dysplasia assessment

The grade of OED was evaluated based on the features recommended by WHO, which includes architectural changes in the surface epithelium and individual cellular alteration during dysplasia. The oral onco-pathologists assessed in total thirty-six biopsy specimens histologically confirmed as OPMDs and graded as mild, moderate, or severe (Fig 1). Dysplastic severity index (SI) was generated for all of them. Features which have been considered for SI calculation according to recommendation of WHO and advice of onco-pathologists are given in Table 1.

**Table 1.** Histological and cytological features of dysplasia used for generating the disease SI.

| **Tissue architectural alterations of oral surface epithelium** | **Individual cellular alterations** |
| --- | --- |
| 1. Bulbous or drop shaped rete ridges | 1. Enlarged nuclei |
| 2. Loss of polarity | 2. Enlarged cells |
| 3. Keratinization of cells below keratinized layer | 3. Increased number and size of nucleoli |
| 4. Loss of epithelial cell cohesion | 4. Increased nuclear cytoplasmic ratio |
| 5. Irregular epithelial stratification | 5. Hyperchromatic nuclei |
| 6. Increased mitotic activity | 6. Pleomorphic nuclei |
| 7. Abnormal mitotic figures | 7. Pleomorphic cells |
| 8. Up to which epithelial layer cellular changes has occurred |  |

Features mentioned in Table 1 are qualitatively evaluated by onco-pathologists in routine diagnosis. In order to appreciate their assessment, we have defined a fuzzy scale in terms of linguistic variables which is described in detail section 2.3.2.

### 2.3 Calculation of severity index

OED grade evaluation is cumulative effect of abnormality of individual features along with their weights. Flowchart of the entire process is shown in Fig.2. The entire procedure may be represented through the following sequential stages, where the first two stages are feature and patient centric information, and the third stage is to provide the Severity Index for each patient to indicate the disease condition of patient.

Stage 1: Determination of aggregated weights of features from different expert’s opinions

Stage 2: Abnormality scoring of individual features in all patients

Stage 3: Calculation of Severity Index of all patients

#### 2.3.1 Stage 1: Determination of aggregated weights of features from different expert’s opinions

Let us consider a group of *n* oral onco-pathologists (hereafter referred as experts) {*E*_1_,*E*_2_,…,*E*_*n*_} and a set of *m* features {*f*_1_,*f*_2_, …,*f*_*m*_} with *m* weights of these given by each of *n* experts. It is also assumed that the experts may have some hesitancy to exercise their options regarding importance of features in OED evaluation. An array of confidence levels of *m* weights is also considered in the system. In this study, eight experts were asked independently to provide weight for each feature given in Table 1 in 0-10 scale (0 being lowest). Hence, *n* = 8 and *m* = 15. Confidence level (scale 1-10, 1 being lowest) shows how much the expert is confident in giving that score.

##### Step 1: Converting crisp weight to fuzzy value by integrating confidence level

Say, for feature *f* an expert assigns weight *w* with degree of confidence *c*. So, the degree of uncertainty viz. *u* = 10 − *c*. Though the weight scale of 10 points is a combination of discrete points, but inclusion of uncertainty makes it a continuous scale. Uncertainty can only be captured through a fuzzy scale, where opting 8 means there is a maximum possibility viz. one of getting 8 but there exists a range around 8 with possibility lower than one. To integrate confidence level with weight, we have utilized the concept of triangular fuzzy number (TFN) here [14]. If an expert is completely uncertain i.e. if *u* = 9, the fuzzy weight 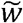 will be a triangular fuzzy number (0,*w*,10). Here, ‘∼’ represents fuzziness. On the other hand, if the expert is completely confident i.e. if *u* = 0, the weight 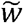 will be (*w,w,w*).

For any other uncertainty level *u*, corresponding fuzzy weight 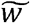 of *f* is:

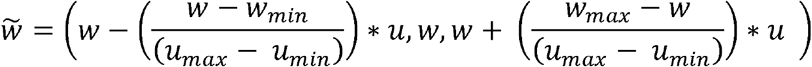

Here, *w*_*min*_ = 0, *w*_*max*_ = 10; *u*_*min*_ =0, *u*_*max*_ = 9, so

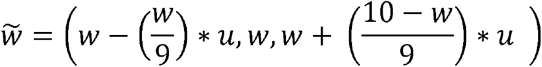

Thus crisp weights of each feature are converted to triangular fuzzy numbers by including the degrees of confidence of the experts.

##### Step 2: Calculation of weight and similarity matrix from experts’ opinions

Let 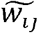 be the fuzzy weight given by *i*^*th*^ expert *E*_*i*_, (*i* =1,2 …,*n*) for the *j*^*th*^ feature *f*_*i*_, (*j* = 1,2, …, *m*). In matrix notation, the weight matrix 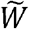, considering all experts’ opinion, against each feature, can be represented as follows:

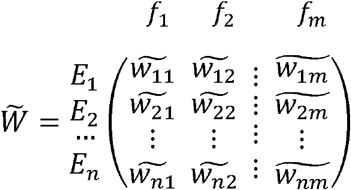

In order to achieve degree of agreement, the calculation of similarity / dissimilarity measure is necessary. If *Ã* ≡ (*a*_1_,*a*_2_,*a*_3_) and 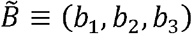 are two triangular fuzzy numbers with corresponding 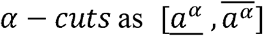 and 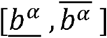 respectively, then distance between two triangular fuzzy numbers *Ã*and 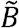 is calculated according to equation given by [15] as follows:

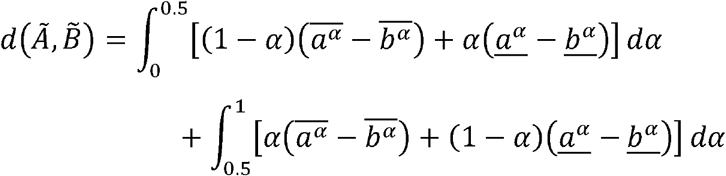

Therefore, similarity measure between two triangular fuzzy numbers *Ã*and 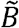 is obtained as 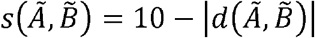. In our case 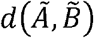 can vary from 0 to 10.

Let *s*_*itj*,_ (1 ≤ *i,t* ≤ *n*) be the similarity measure between the pair of experts’ *E*_*i*_ and *E*_*t*_ opinions against *j*_*th*_ feature, then a similarity matrix *AM*_*j*_ against *j*_*th*_ feature is considered as follows.

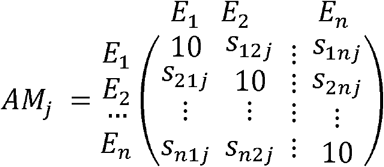

Here, the agreement between the opinions of the expert *E*_*i*_ and *E*_*t*_ for *j*^*th*^ feature is 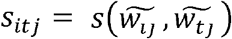 for *i* ≠ *t* and *s*_*itj*_ = 10 for *i* =*t*. Therefore, *m* number of *n* ×*n* similarity matrices or agreement matrices will be achieved.

##### Step 3: Calculation of the average agreement degree and relative agreement degree of each expert against each feature

For each feature *f*_*j*_(*j* = 1,2..*m*), the average agreement degree of any expert *Ei* (*i* = 1,2,..*n*) is defined in following way,

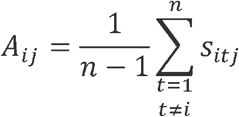

Overall agreement against each feature is calculated as

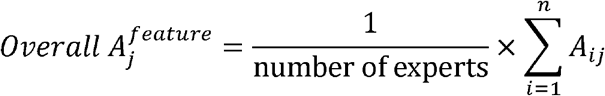

If all the experts are in perfect harmony or unanimous in opinions, the overall agreement against each feature is ten.

The relative agreement degree *R*_*ij*_ of *i*^*th*^ expert for *j*^*th*^ feature,

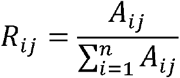

which ensures the fact that when all the experts are unanimous in opinions, each relative agreement degree is 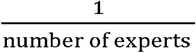.

##### Step 4: Calculation of overall weight of each feature

Finally, for each feature *f*_*j*_(*j* = 1,2..*m*), an aggregated weight, *Aw*_*j*_, considering all experts is defined as follows:

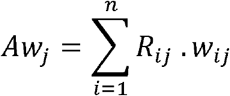

Aggregated weight of each feature is then normalized to get sum of weights of all features equal to 1. Final normalized aggregated weight of *j*^*th*^ feature 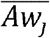 is given by

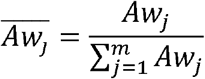

Supplementary section A shows an example of aggregated weight calculation procedure for a single feature.

#### 2.3.2 Stage 2: Abnormality scoring of each feature in individual patient

Onco-pathologists evaluated abnormality of all features of Table 1 using linguistic variables instead of numerical values as experts are more comfortable to express their opinion in linguistic terms [16]. The theory of fuzzy logic enables us to deal with such general sources of vagueness [14, 17, 18]. A general linguistic scale, ranging from normal to most extreme diseased conditions, may be considered to capture experts’ opinion in this regard. Features may not exhibit any change from normal or the change can be very low to very high. This scale (Table 2a) may be used for all features mentioned in Table 1 except three tissue architectural features viz. loss of epithelial cell cohesion (LECH), irregular epithelial stratification (IES) and up to which epithelial layer cellular changes has occurred (ELC). These three features are noted as per their association with epithelial layer i.e. expression of these features till basal-supra basal or lower third or middle third or upper third or full thickness of epithelium (Table 2b). Both Table 2a and 2b indicate abnormality score of each feature in terms of generalized triangular fuzzy number (TFN).

**Table 2a.** General fuzzy scale for linguistic evaluation of features’ abnormality.

| Linguistic variables w.r.t. normal and corresponding TFN |  |
| --- | --- |
| No change/absent | (0, 0, 0) |
| Very low | (0, 0, .25) |
| Low | (0, .25, .5) |
| Medium | (.25, .5, .75) |
| High | (.5, .75, 1) |
| Very high | (.75, 1, 1) |

**Table 2b.** Fuzzy scale for features expressed in terms of association with epithelial layer.

| Epithelial layer till which LECH, IES and ELC are present and corresponding TFN |  |
| --- | --- |
| Absent | (0, 0, 0) |
| Basal-supra basal | (0, 0, .25) |
| Lower third of epithelium | (0, .25, .5) |
| Middle third of epithelium | (.25, .5, .75) |
| Upper third of epithelium | (.5, .75, 1) |
| Full thickness of epithelium | (.75, 1, 1) |

For increased number and size of nucleoli, hyperchromatic nuclei and pleomorphic nuclei (cells), linguistic variables (Table 2a) correspond to percentage of nuclei (cells) undergone change in a region of interest. Very low, low, medium, high and very high correspond to less than 20%, 20-40%, 40-60%, 60-80% and greater than 80% except increased number and size of nucleoli. For increased number and size of nucleoli, very low, low, medium, high and very high correspond to less than 15%, 15-30%, 30-45%, 45-60% and 60-75%.

Further, enlarged nuclei, enlarged cells, and increased nuclear-cytoplasmic ratio are layer wise evaluated to increase the accuracy of OED grading. Severity of OED increases along with progression of cellular changes to the upper layers [19]. Following weights have been used for different layers: 0.1 for basal-supra basal, 0.2 for lower third of epithelium, 0.3 for middle third of epithelium and 0.4 for upper third of epithelium. These weights are chosen as sum of weights of all layers should be 1 otherwise abnormality score (TFN) of these three features may be greater than (1, 1, 1) as can be seen from examples in supplementary (section B). Both percentage of cell or nuclei which have undergone morphometric change in a region of interest and their degree of changes are considered in each layer. For degree of change, nuclei and cells are qualitatively compared with relatively normal region of same tissue. Both attributes i.e. percentage and degree are evaluated by general fuzzy scale of Table 2a viz. very low to very high. For degree of change in nuclear and cell size, very low, low, medium, high and very high correspond to less than 15%, 15-30%, 30-45%, 45-60% and 60-75% of original size. Average of these two components (viz. percentage and degree of change) is calculated for each layer as triangular fuzzy number cannot be greater than (1, 1, 1). This average value is multiplied by corresponding layer weight and all the layers are added to obtain abnormality score of these three features. Supplementary (section B) shows two examples of abnormality scoring for enlarged nuclei.

#### 2.3.3 Calculation of severity index

Abnormality score of each feature (TFNs of Table 2a and 2b) is multiplied with weight of that feature and all the features are added to get fuzzy severity index 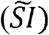.

Let, abnormality score of feature *f*_*j*_(*j* = 1,2..*m*) be 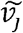 and its weight be 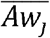.

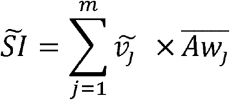

Centroid based defuzzification method [20] is done to convert fuzzy SI to a crisp valued index *SI*. *SI* can vary between 0 and 1.

## 3. Results and discussion

During the evaluation of degree of severity for any disease, physicians add different mental weightage to different parameters (clinical features, histopathological features, patient’s previous medical history, etc.) and sub-parameters before making a final diagnostic assessment — which directly corresponds to treatment plan and prognosis of the disease. Because of the complex nature of the diagnostic assessment, development of algorithm-assisted decision support system is beneficial. Keeping track of all the features simultaneously in diagnostic decision making may account in part for lack of reproducibility which can be improved by an index. Literature review shows such indices exist for other diseases [21-34]. But most of them consider all contributing factors with equal importance. While in few indices, weights are assigned to different contributing attributes according to their clinical significance, these weights are subjective. This study aims to formulate an index after reducing subjectivity of weights of dysplastic features in the context of OED.

Based on their experience and knowledge, oral onco-pathologists vary in their opinion about importance (weights) of features in the context of severity assessment of epithelial dysplasia. In our work we have also considered confidence level of experts which is very important in any real world decision making problem. After collecting weights from eight experts separately, aggregated weights of features as obtained by group decision making algorithm described in section 2.3.1 are shown in Table 3.

**Table 3.** Weights given to each feature using group decision making algorithm.

| Tissue architectural features |  | Individual cellular alterations |  |
| --- | --- | --- | --- |
| Features | Weight | Features | Weight |
| 1. Bulbous or drop shaped rete ridges | 0.054 | 1. Enlarged nuclei | 0.054 |
| 2. Loss of polarity | 0.069 | 2. Enlarged cells | 0.049 |
| 3. Keratinization of cells below keratinized layer | 0.053 | 3. Increased number and size of nucleoli | 0.063 |
| 4. Loss of epithelial cell cohesion | 0.077 | 4. Increased nuclear cytoplasmic ratio | 0.069 |
| 5. Irregular epithelial stratification | 0.065 | 5. Hyperchromatic nuclei | 0.072 |
| 6. Increased mitotic activity | 0.088 | 6. Pleomorphic nuclei | 0.073 |
| 7. Abnormal mitotic features | 0.09 | 7. Pleomorphic cells | 0.065 |
| 8. Up to which epithelial layer cellular changes has occurred | 0.069 |  |  |

SI was calculated for each of the thirty-six slides of histopathologically diagnosed dysplastic patients according to the method described in section 2.3. Table 4 shows p values from Student’s t test confirming distinct values of SI for mild, moderate and severe cases. Fig 3 shows boxplot of SI values for different categories. One patient diagnosed with moderate dysplasia is shown as outlier in boxplot with SI of 0.42. Its’ high SI can be attributed to high mitotic activity, nuclei pleomorphism and marked cellular changes despite cellular alterations being present up to middle third layer of epithelium. For each category, SI can have range of values further indicating low and high risk of individual grade.

**Table 4.** p values of severity index values from two tailed t test @ 1% significance level.

| Mild vs Moderate | Moderate vs Severe |
| --- | --- |
| 5.57E-09 | 1.23E-05 |

**Fig. 2.**
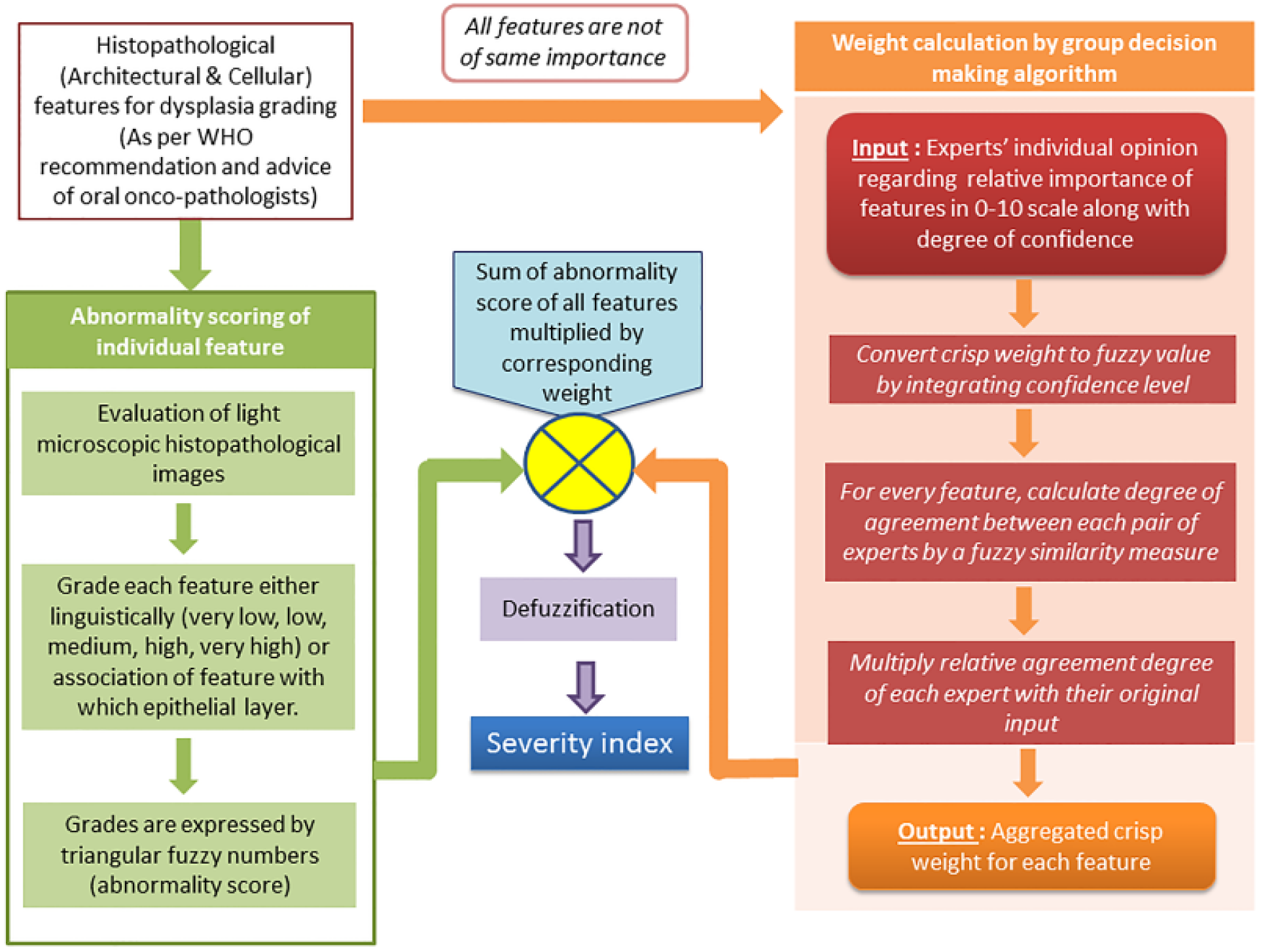
Flowchart for calculating severity index

**Fig. 3.**
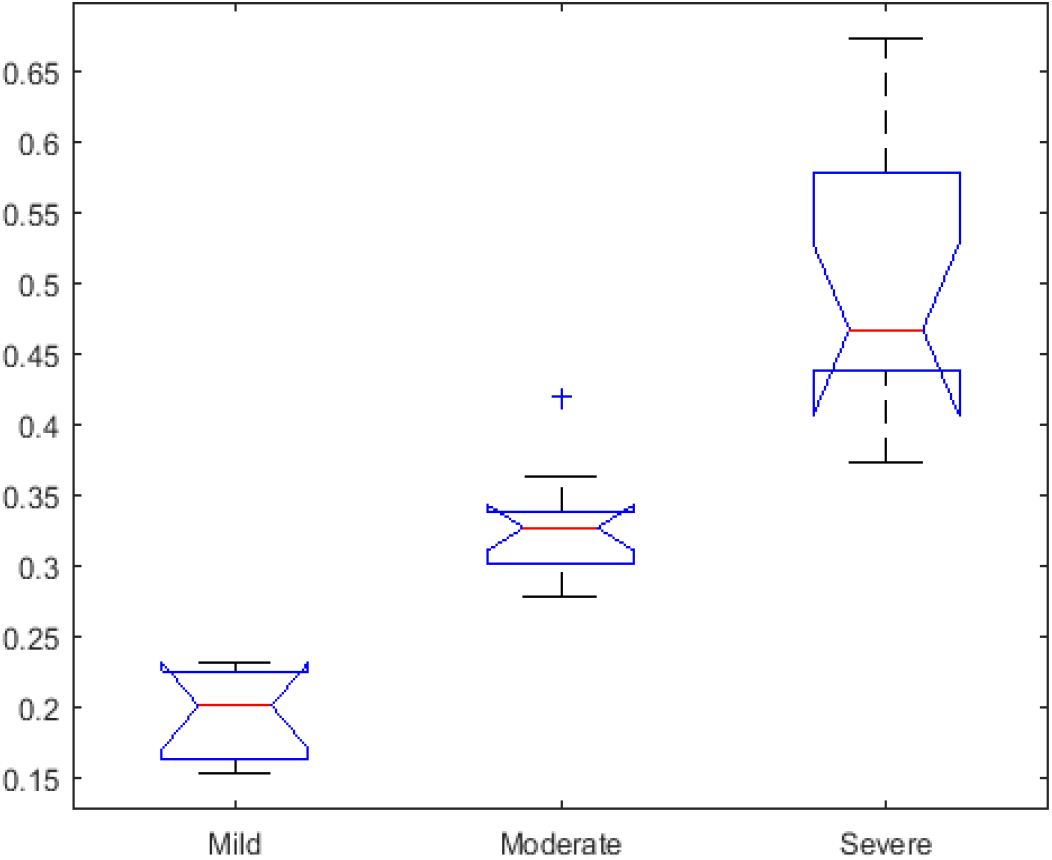
Boxplots showing severity index values for different grades of OED.

To illustrate the applicability of SI, histopathological data of five patients (P1, P2, P3, P4 and P5) are shown in Table 5a. P1 belongs to mild, P2 and P3 belong to moderate and P4 and P5 belong to severe dysplasia category. Table 5b shows calculated SI values of these five patients. It can be noted that mitotic activity and abnormal mitotic figures, two most important features are more in P2 than P3. But considering all features SI is higher for P3 than P2 as shown in Table 5b.

**Table 5a.** Evaluation of five patients by oral onco-pathologist.

| Feature | P1<br>(mild) | P2<br>(mod) | P3<br>(mod) | P4<br>(severe) | P5<br>(severe) |
| --- | --- | --- | --- | --- | --- |
| Bulbous or drop shaped rete ridges | A | A | A | B | A |
| Loss of polarity | VL | M | H | H | VH |
| Keratinization of cells below keratinized layer | A | A | A | A | A |
| Loss of epithelial cell cohesion | B-SB | LT | MT | MT | UT |
| Irregular epithelial stratification | LT | LT | MT | MT | UT |
| Increased mitotic activity | H | M | L | VL | VH |
| Abnormal mitotic figures/abnormal superficial mitosis | VL | L | A | VL | M |
| Epithelial layer till which cellular changes have occurred | LT | MT | MT | UT | UT |
| Enlarged nuclei | Till LT:<br>H%+M | Till MT:<br>M%+L | Till MT:<br>H%+M | Till MT:<br>H%+H<br>and in<br>UT:<br>H%+M | Till UT:<br>VH%+<br>H |
| Enlarged cells | Till LT:<br>M%+M | Till MT:<br>M%+L | Till MT:<br>H%+L | Till UT:<br>H%+M | Till UT:<br>VH%+<br>M |
| Increased number and size of nucleoli | L | VL | L | VL | H |
| Increased nuclear cytoplasmic ratio | In B-SB:<br>H%+M<br>and in<br>LT:<br>L%+M | Till MT:<br>M%+L | Till MT:<br>H%+M | Till UT:<br>H%+M | Till UT:<br>VH%+<br>H |
| Hyperchromatic nuclei | M | M | H | H | L |
| Pleomorphic nuclei | L | L | M | M | H |
| Pleomorphic cells | VL | L | L | L | H |

[mod: moderate; A: absent; B: bulbous; VL: very low, L: low, M: medium, H: high, VH: very high; B-SB: basal-supra basal, LT: lower third, MT: middle third, UT: upper third of epithelium]

**Table 5b.** Computed severity index (SI)

| Patient | Diagnosis | SI |
| --- | --- | --- |
| P1 | Mild | 0.227 |
| P2 | Moderate | 0.285 |
| P3 | Moderate | 0.363 |
| P4 | Severe | 0.44 |
| P5 | Severe | 0.636 |

## 4. Conclusion

A fuzzy logic-based group decision making algorithm has been used for weights calculation of different histopathological and cellular parameters to reduce disparity in assessment of OED grading. Inclusion of confidence level of experts makes the process of aggregated weight generation more robust by mimicking real life situation. Advancement of soft computing tools prompts us to consider the subjective opinions with proper stress. For abnormality grading of individual feature, fuzzy logic, known for its ability to capture subjective input with vagueness is employed. The overall aim of our study is to assist oncologists’ decision-making regarding severity grading of OED. To assess combined effect of the attributes used in the OED grading in a more reproducible way, we have proposed SI. SI will help to stratify patients belong to same dysplasia grade in low or high end of that grade. Proposed SI will help in choosing more effective and appropriate therapeutic strategies for OED and consequently help in achieving better prognosis. Present methodology may be applied to other medical decision making problems and for developing indices of various other diseases.

## Acknowledgment

The authors acknowledge SMST and ATDC dept., Indian Institute of Technology (IIT) Kharagpur, Guru Nanak Institute of Dental Science and Research (Kolkata) for providing research facility and IOP project (IIT/SRIC/MM/IOP/2017-18/208) of Higher education, (Sci and tech), Govt of West Bengal for research funding.

