## Supplementary material for "Development of Severity Index for Oral Epithelial Dysplasia Using Fuzzy Group Decision Making Algorithm": Tables

**A. An example of aggregated weight calculation procedure for any feature**

This section shows an example of aggregated weight calculation for any feature as described in section 2.3.1. Weights given by eight experts and their confidence levels are shown in Table A.1.

**Table A.1** Weights given for a particular feature by eight experts along with their confidence level

|  | **Expert 1** | **Expert 2** | **Expert 3** | **Expert 4** | **Expert 5** | **Expert 6** | **Expert 7** | **Expert 8** |
| --- | --- | --- | --- | --- | --- | --- | --- | --- |
| **Weight** | 5 | 3 | 8 | 7 | 3 | 6 | 8 | 5 |
| **Confidence level** | 10 | 7 | 6 | 8 | 6 | 9 | 7 | 6 |

1st expert has assigned weight 5 with confidence level 10. 2nd expert has assigned weight 3 with confidence level 7. Next, these weights are transformed to fuzzy value (Table A.2) by integrating confidence level.

**Table A.2** Fuzzified weight of eight experts by integrating confidence level

|  | **Expert 1** | **Expert 2** | **Expert 3** | **Expert 4** | **Expert 5** | **Expert 6** | **Expert 7** | **Expert 8** |
| --- | --- | --- | --- | --- | --- | --- | --- | --- |
| **Fuzzified**  **weight** | (5, 5, 5) | (2, 3, 5.33) | (4.44, 8, 8.89) | (5.44, 7, 7.67) | (1.67, 3, 6.11) | (5.33, 6, 6.44) | (5.33, 8, 8.67) | (2.78, 5, 7.22) |

Table A.3 shows similarity metric i.e. degree of agreement between each pair of experts. Corresponding average agreement degree and relative agreement degree for each expert are also shown.

**Table A.3** Pairwise similarity between experts and individual agreement degree

| Similarity between each pair of experts | | | | | | | | |  |  |  |
| --- | --- | --- | --- | --- | --- | --- | --- | --- | --- | --- | --- |
|  | **Expert 1** | **Expert 2** | **Expert 3** | **Expert 4** | **Expert 5** | **Expert 6** | **Expert 7** | **Expert 8** |  | **Average**  **agreement**  **degree** | **Relative agreement degree** |
| **Expert 1** | 10 | 8.75 | 7.11 | 7.94 | 9 | 8.92 | 7.08 | 9.44 |  | 8.32 | 0.13 |
| **Expert 2** | 8.75 | 10 | 5.86 | 6.69 | 9.75 | 7.67 | 5.83 | 8.19 |  | 7.54 | 0.117 |
| **Expert 3** | 7.11 | 5.86 | 10 | 9.17 | 6.11 | 8.19 | 9.97 | 7.67 |  | 7.73 | 0.12 |
| **Expert 4** | 7.94 | 6.69 | 9.17 | 10 | 6.94 | 9.03 | 9.14 | 8.5 |  | 8.2 | 0.128 |
| **Expert 5** | 9 | 9.75 | 6.11 | 6.94 | 10 | 7.92 | 6.08 | 8.44 |  | 7.75 | 0.121 |
| **Expert 6** | 8.92 | 7.67 | 8.19 | 9.03 | 7.92 | 10 | 8.17 | 9.47 |  | 8.48 | 0.132 |
| **Expert 7** | 7.08 | 5.83 | 9.97 | 9.14 | 6.08 | 8.17 | 10 | 7.64 |  | 7.7 | 0.12 |
| **Expert 8** | 9.44 | 8.19 | 7.67 | 8.5 | 8.44 | 9.47 | 7.64 | 10 |  | 8.48 | 0.132 |

Aggregated weight= 0.13*5+0.117*3+0.12*8+0.128*7+0.121*3+0.132*6+0.12*8+0.132*5=5.63

**B. Abnormality scoring of enlarged nuclei, enlarged cells, and increased nuclear-cytoplasmic ratio in individual patient**

Following two examples show method of scoring enlarged nuclei. Same process is followed for enlarged cells and increased nuclear cytoplasmic ratio.

a) Say, high percentages of nuclei are enlarged by medium amount till lower third of epithelium. In this case, for basal layer abnormality of nuclei will be 0.1$\times$0.5$\times$ (High% + Medium) and for lower third of epithelium abnormality of nuclei will be 0.2$\times$0.5$\times$ (High% + Medium).

Final abnormality score for nuclei will be

0.1$\times$0.5$\times$ (High% + Medium) + 0.2$\times$0.5$\times$ (High% + Medium) = 0.3$\times$0.5$\times$ (High% + Medium) = 0.3$\times$0.5$\times$ ((0.5, 0.75, 1) + (0.25, 0.5, 0.75)) =0.3$\times$0.5$\times$ (0.75, 1.25, 1.75)

b) Nuclei are enlarged till upper third of epithelium. But up to middle third, high percentages of nuclei are enlarged and amount of enlargement is medium where as in upper third, medium percentages of nuclei are enlarged and amount of enlargement is medium. In this case severity will be

0.6$\times$0.5$\times$ (High% + Medium) + 0.4$\times$0.5$\times$ (Medium % + Medium)

=0.6$\times$0.5$\times$ ((0.5, 0.75, 1) + (0.25, 0.5, 0.75)) +0.4$\times$0.5$\times$ ((0.25, 0.5, 0.75) + (0.25, 0.5, 0.75)).
